# Passive visual stimulation induces fatigue or improvement depending on cognitive load

**DOI:** 10.1101/2020.11.19.390096

**Authors:** Stefano Ioannucci, Guillermo Borragán, Alexandre Zénon

## Abstract

Theories of mental fatigue disagree on whether performance decrement is caused by motivational or functional alterations. We tested the assumption that keeping neural networks active for an extensive period of time entrains consequences at the subjective and objective level – the defining characteristics of fatigue – when confounds such as motivation, boredom and level of skill are controlled. We reveal that passive visual stimulation affects the performance of a subsequent task that is carried out in the same portion of visual space. This outcome, consisting either in an enhancement or deterioration of performance, was determined by the participants’ level of cognitive load and arousal, which were manipulated through variations in the difficulty of concurrent auditory tasks. Thus, repeated stimulation of neural networks leads to their altered functional performance, a mechanism which may play a role in the development of global mental fatigue.

## Introduction

Prolonged mental activity leads to a sensation of discomfort with well-known adverse consequences on overall quality of life (Åkerstedt et al., 2007; Raslear et al., 2011), productivity (Ricci et al., 2007), impulsivity (Blain et al., 2016), emotional regulation (Grillon et al., 2015) and workplace accidents (Goode, 2003; Swaen et al., 2003), all regrouped under the umbrella term of Mental Fatigue (MF). MF manifests itself in two empirically measurable components: the *subjective* feeling of tiredness and the *objective* decrease in performance (DeLuca, 2007). To this day, the neural origin of MF remains elusive, and this lack of understanding stems in part from MF’s dependence on the duration and difficulty of the task (Hockey, 2013), as well as on the motivation (Boksem et al., 2006), fatiguability and skills of the agent performing it (Borragán et al., 2017).

Theories of mental fatigue generally concur on its role in disengaging individuals from cognitive tasks to seek rest. Yet, the underlying cause of this need to withdraw from effortful tasks continues to be debated by two schools of thought. Functional theories assume that fatigue arises as a consequence of alterations in the neural circuitry (e.g. depletion of finite resources or accumulation of metabolites), or as an outcome of mechanisms that prevent such alterations (Blain et al., 2016; Dongen et al., 2011; Schellekens et al., 2000; Zénon et al., 2019). Despite their intuitive appeal, these accounts have suffered from a lack of evidence (Hockey, 2013; Kurzban et al., 2013). In response, motivational theories view fatigue as a cognitive strategy, responsible for the reallocation of effort when the cost-benefit ratio of ongoing behaviour is disadvantageous (Boksem & Tops, 2008; Hopstaken, Linden, et al., 2015; Kurzban et al., 2013). However, this motivational view of fatigue has also been challenged (Benoit et al., 2019; Gergelyfi et al., 2015).

From a neuroscientific perspective, the fact that fatigue may arise from any type of task complicates the process of pinpointing it to specific brain areas (Harrington, 2012), due to the risk of conflating fatigue-specific effects with unrelated activity across brain regions. Rigorous exploration of fatigue should first contemplate its functioning at the lowest degrees of complexity, to rule out as many confounds as possible. A prediction of the functional paradigm is that repeated stimulation of specific neural networks should lead to their altered performance over time.

We coin as *neural fatigue* this putative specific functional deterioration, which we tested by drawing inspiration from a line of research on sleep that found a decrease in visual perception performance due to repetitions of a visual task, the texture discrimination task (TDT), where participants are required to identify the orientation of briefly presented peripheral targets (Mednick et al., 2002). Remarkably, in this study, participants’ performance was restored when perceptual characteristics of the targets (orientation and location) were changed, but not when they were offered money to manipulate motivation. Additionally, a follow-up neuroimaging study found this perceptual deterioration to correlate with decreased BOLD signal in V1 (Mednick et al., 2008). Here, we sought to determine if this perceptual deterioration in the TDT task could be passively induced by prolonged stimulation of specific portions of the visual field.

Given gain theories of arousal (Aston-Jones & Cohen, 2005; Mather et al., 2016), we predicted that being engaged in concomitant auditory tasks of varying cognitive load during saturation would modulate its impact on behaviour. Our hypothesis was to expect a stronger effect in agents performing tasks with harder demands, given the stimulating effects of arousal on cortical response to stimulation (Jones, 2003; Mather et al., 2016; McGinley et al., 2015). Furthermore, cerebral activity was recorded by electroencephalography (EEG) to assess alterations in the steady-state response at stimulation frequency (∼7.5 hertz), as we were expecting to disrupt low-level bottom-up visual processing. Lastly, the relationship between behavioural consequences of saturation and subjective feeling of fatigue was investigated.

## Methods

Forty-eight participants took part voluntarily in the experiment (24m and f, age = 22.1 ± 2.24). All of them were naive to the experimental procedure, with no history of mental or visual conditions. Participants had normal or corrected-to-normal vision and provided written informed consent to participate. They received 10€ per hour in compensation for their participation. The study was approved by the Ethical Review Board. Due to the lack of literature on the effect we sought to test, the sample size was arbitrarily preplanned to have a balanced and sufficient number of participants for the various experimental conditions and randomisations.

In total, 56 participants were recruited, given that 7 participants were excluded; 4 due to failure to respond to an adequate number of trials in the behavioural task (less than half the trials in two or more blocks of the first session), 2 spontaneously dropped out during the experiment and 1 for being an outlier (performance > 4 std).

### Experimental design

All aspects of the experiment, except the questionnaires, were implemented through Matlab 2019a (The MathWorks, Inc., Natick, Massachusetts, United States), using Psychtoolbox (Brainard, 1997; Pelli, 1997) and displayed on a computer screen with 1280 by 1024 resolution. Participants sat in front of the screen at a distance of 60 cm and accommodated their heads on a chin-rest. A keyboard was used to respond across the various tasks. Below the screen there was an Eyetracker 1000 (SR Research Ltd., Mississauga, Canada), which was employed to ensure central fixation of the participants’ gaze while tracking variations in the size of their pupils. A 32-channel Active Two Biosemi system (Biosemi, Amsterdam, Netherlands) EEG headset was mounted on their heads. Lastly, a pair of in-ear headphones was provided (Cellularline spa., Reggio Emilia, Italy).

The experimental design was divided into different phases, which will be detailed in the following sections. First of all, a brief training with reduced number of trials of the various tasks was carried out in order to have the participants familiarise with their procedures (see Figure 1 panel 1a). Behavioural and physiological measures were taken at different timepoints of the experiment, namely: baseline, middle and conclusion (see Figure 1 panel 1b,d,f). In-between these measurements, subjects underwent continuous stimulation (‘saturation’) of a specific portion of their visual field (‘quadrant’) (see Figure 1 panel 1c,e). During saturation, volunteers were engaged in auditory tasks, the difficulty of which differed based on the experimental group to which they were randomly assigned, being either in an “easy” (n=24, 12m and f, age: 22.37 ± 2.7) or “hard“ (n=24, 12 m and f, age: 21.92 ± 1.69) condition. The assignment of saturated quadrant and group was counter-balanced across participants. Overall, an experimental run lasted approximately 2h and 30 minutes, depending on the time spent to install the EEG headset.

**Figure 1:**
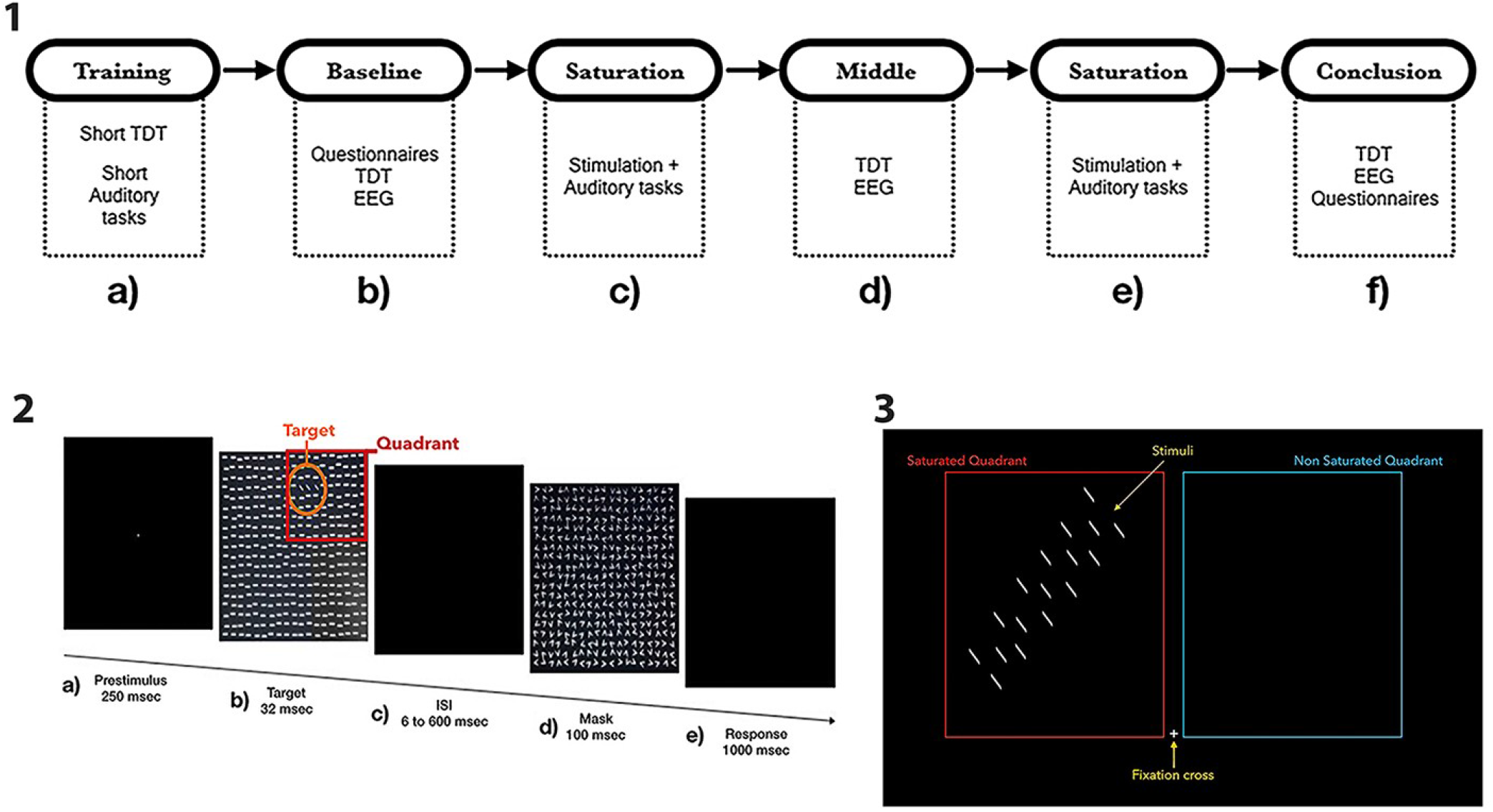
Panel 1: Flowchart of the experimental design. Panel 2: Visual representation of the Texture Discrimination Task (TDT). Please note that across the whole trial a fixation cross stayed in the centre in order to help participants maintain their gaze fixated. Panel 3: Visual representation of saturation and EEG sessions. While fixating the centre a participant would be presented with stimuli identical to the targets of the TDT (here they appear larger due to representation), appearing in all possible target locations within a quadrant at 7.5 Hz. During EEG sessions the quadrants alternated (see EEG methods for details), while during saturation sessions the stimuli flashed in a single quadrant (the saturated quadrant), and participants were engaged in concurrent auditory tasks, the difficulty of which depended upon their experimental group

### Auditory tasks

As outlined, during each saturation session participants were engaged in auditory tasks, the demands of which varied according to the experimental group to which they were assigned for the duration of the whole experiment, being either easy or hard. This experimental condition was introduced to evaluate potential effects of differing cognitive loads, and resulting arousal levels, on the time course of fatigue. Therefore 3 different tasks were adopted, based on the fact that their complexity could be manipulated and that they provided variety in demands (working memory, executive functions, attention). These tasks were randomly cycled across a saturation session, until its pre-scripted duration of 41 minutes had passed. Specifically, the following tasks were used: the n-back task, a pitch-sequence reproduction task, and a switch task which we named the side task (Table 1). All tasks of the experiment employed the same response keys (F and J), except for the pitch-sequence task where 2 additional keys were added (D and K).

**Table 1:** Summary of the demands in the auditory tasks (rows) by difficulty condition (columns)

| TASK | EASY GROUP | HARD GROUP |
| --- | --- | --- |
| N-back | Press key when hearing the letter X | Press key when current letter corresponds to a letter presented 2 steps before in the sequence |
| Pitch-sequence | Reproduce a sequence of two beeps | Reproduce a sequence of 5 to 8 faster beeps |
| Side | Indicate if a sound was played through the left or the right headphone | Same as easy, but the response keys could be inverted throughout the block depending on sound category. Inhibit response to an additional category |

N-back: a trial would consist of a list of 12 letters where participants had to report the occurrence of the target letter. In the easy group, the target letter was ‘X’ (0-back), while in the hard group the target letter was any letter repeated 2 steps before (2-back). A block of N-back was composed of 35 trials for both conditions, each trial lasted 15 seconds for both groups.

Pitch-sequence: In this task, participants were required to replicate a sequence of beeps that was presented to them. Four different beeps (from low-pitch (336hz) to high-pitch (475hz) and in between (377hz, 424hz) were put together in randomly generated sequences. To each beep corresponded a key on the keyboard. For the easy group these sequences comprised 2 beeps, while for the hard group they comprised 5 to 8 beeps. A block of pitch-sequence task consisted in 80 trials for both conditions, with longer intervals between beeps for the easy group and shorter intervals for the hard group. On average, the duration of stimuli presentation for the easy group lasted 2.7 seconds, while the mean duration for the hard group was of 8.9 seconds.

Side: In this task sounds were presented randomly either to the left or to the right earphone of the participant. These sounds came from different categories, namely: animal sounds or vehicle sounds. For the easy group, participants only had to indicate if the current sound was played on the left or the right. The hard group had instead a cue voice indicating, at random points during the block, which category they had to answer coherently to (i.e. if a sound of that category was presented to the left, they had to press the left key and *vice-versa*), this implicitly signalled they had to answer incoherently to the unmentioned category (i.e. if the sound was on the left, they had to press right and *vice-versa*). Furthermore, for this group a third category of sounds was added, the computer/electronics category, to which they were instructed not to respond. A block would be made up of 135 sounds with shorter silences in-between for the hard group, and 80 sounds with longer distances in-between for the easy group, subdivided into trials of 5 sounds for both groups. A trial would last on average 30 seconds in the easy condition, and 18 for the hard group.

### Behavioural index of fatigue

As a behavioural measure of fatigue, we adopted the participants’ performance in the Texture Discrimination Task (TDT), based on the task developed by Karni and Sagi (Karni & Sagi, 1991). The task’s goal is to discriminate the orientation of a peripheral target, which consists of three diagonal lines aligned either vertically or horizontally, against a background of horizontally oriented bars (see Figure 1 panel 2b). Participants were instructed to maintain their gaze on a central fixation cross and report the perceived orientation of a peripheral target at the end of each trial. These targets would relocate from trial to trial, within a defined quadrant of the screen. Quadrants would vary depending on the block, either on the upper right or upper left portion of the screen. Each experimental trial comprised a pre-stimulus window of 700 ms (see Figure 1 panel 2a), followed by the target screen for 32 ms (see Figure 1 panel 2b), a blank screen, known as ISI (inter stimulus interval), with variable length between 6 and 600 ms, determined by a Bayesian staircase procedure (see below) (see Figure 1 panel 2c), followed by a mask of 100 ms (see Figure 1 panel 2d). The mask is designed to disrupt the subjects’ processing of the peripheral target lines, therefore shorter ISIs translated into harder trials.

Finally, there was a 1000 ms time window where the participant indicated, by pressing on the keyboard, if the target was perceived as horizontal or vertical (see Figure 1 panel 2e). A single TDT trial lasted on average 2.26 seconds (±0.28 sec) and 80 trials composed a block. A complete TDT session consisted of 4 blocks, 2 per quadrant, and lasted 12.4 minutes on average (± 37 sec). The order of blocks was random for every session and participant. A block would begin when the participant pressed spacebar, after which he or she was informed on the quadrant within which the target would appear, by means of a message at the centre of the screen.

### Adaptive ISI selection procedure

In order to optimize inference of participants’ psychometric curve from limited data, we opted for an adaptive procedure to select ISI values (inspired from (Kontsevich & Tyler, 1999)). This procedure was applied following the five first trials (in which ISI was selected randomly). In each trial, a variational Bayesian logistic regression was performed on currently acquired data (Drugowitsch, 2019), resulting in parameter estimates and variance. Then, the expected update in these parameters was computed under the hypotheses of correct or incorrect response in the next trial, and for all possible ISI values:

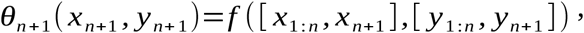

for *θ* the mean (*θ^μ^)* and variance (*θ^Σ^)* estimates of the parameters, *f* the variational logistic regression, *y*_*1:n*_ the acquired response data and *y*_*n+1*_ the expected response, *x*_*1:n*_ the past ISI values and *x*_*n+1*_ the ISI value for the next trial.

The Kullback-Leibler (KL) divergence between current and future estimates, representing how much information one would expect to gain about the parameter estimates, was computed for each ISI and possible response according to the following formula:

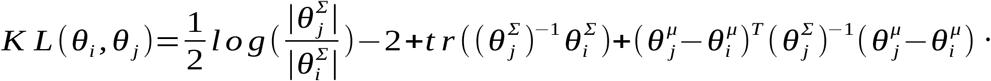

These KL values for correct and incorrect expected response were then averaged and weighted as a function of the probability of obtaining either correct or incorrect response given current parameter values:

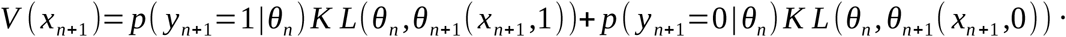

These KL values were then softmax-transformed to lead to a probability distribution over all ISI values, which was then sampled randomly to select the value of the ISI for the next trial.

### Saturation

A saturation session lasted for 41 minutes and consisted of a prolonged visual stimulation, during which participants had to maintain their gaze on a central fixation point while stimuli were continuously flashed at 7.5Hz (see Figure 1 panel 3). Stimuli were composed of all the possible targets within a quadrant in the TDT. The saturated quadrant was kept constant for each participant across all saturation sessions. This ensured the entraining of specific neuronal populations across the whole experiment, providing a way to test whether the neural (EEG) and behavioural (TDT) response would differ between saturated and non-saturated quadrants.

### EEG

EEG recordings of whole brain activity in response to the presentation of the saturation stimulus (See Figure 1 panel 3) were performed. A complete EEG session comprised four repetitions of 1-minute long stimulations (two per quadrant), with a 5-second pause in between. Visual stimulation during EEG blocks was identical to the one used during saturation blocks, except that the quadrants with stimuli alternated, their order being counterbalanced across participants.

FieldTrip toolbox (Oostenveld et al., 2011) was employed for the analysis of EEG data. Specifically, we focused our analysis on the stimulation frequency (7.5 Hz), reasoning that alterations in bottom-up visual processing should lead to modulation of the steady-state response.

The preprocessing consisted in a low-pass filter at 100 hertz, a high-pass filter at 0.4 hertz and a line-noise filter at 50 and 100 hertz applied to the raw data, which was then re-referenced on the average of all 32 channels. Before segmentation into trials, artefacts were rejected upon visual inspection of the data and then by independent component analysis. Subsequently, one Fourier transform with a single Hanning taper was applied to extract the power spectrum around stimulation frequency, (7.3 to 7.7 hertz, in steps of 0.1). Signal from neighbouring frequencies (7.3 to 7.4 and 7.6 to 7.7 hz) was subtracted from the 7.5 Hz signal, to highlight the specific response to stimulation (Norcia et al., 2015), and this noise-subtracted 7.5hz was then used in subsequent analyses.

### Pupil

We also recorded the pupil size of participants as a physiological index of arousal, as it is a well-established physiological marker of this construct (McGinley et al., 2015; Wang et al., 2018). Variations in pupil size were tracked during saturation sessions, while participants were engaged in the concurrent auditory tasks, the difficulty of which varied according to the experimental group they were assigned to. Specifically, after linear interpolation of blinks, filtering (high-pass with 0.01Hz cutoff frequency) and baseline correction, the pupil responses between the onset of the first stimulus of trial n and the onset of the first stimulus in trial n+1 were averaged. These estimates of trial-wise average pupil responses were then averaged by auditory task and session, thus resulting in one data point for each subject and task in both saturation sessions.

### Questionnaires

In order to measure the participants’ subjective feeling of fatigue and sleepiness, two pen-and-paper questionnaires were administered at the beginning and end of the experiment; the Multidimensional Fatigue Inventory (Gentile et al., 2003) and the Karolinska Sleepiness Scale (Shahid et al., 2011).

### Statistical Analysis

First, behavioural effects were assessed by means of a Generalised Linear Mixed Model (GLMM) on response accuracy in the TDT task. Correct response in the task was modelled as a logistic dependent variable, with ISI as a covariate and participants as a random variable. Difficulty group (easy, hard), experimental session (baseline, middle, conclusion) and saturated quadrant (yes, no) were set as explanatory variables, testing their main effects and interactions. This was done on Jamovi (The jamovi project, 2019) through its module GAMLj (Gallucci, 2019). Random effects were included step-by-step as long as the Bayesian Information Criterion and deviance values kept decreasing and no convergence issues were encountered. Specifically, intercept, ISI and Session were included in the random part of the model and were therefore allowed to vary across participants.

Second, effects on brain activity were evaluated by examining changes in the EEG signal. To evaluate the interaction of session and condition, the difference of EEG activity between saturated and non-saturated condition was computed, for each of the three sessions and for each participant.

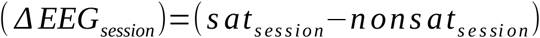

These values were then analysed in dependent sample two-tailed t-tests to assess changes across sessions. Correction for multiple comparisons across electrodes was enforced by means of a non-parametric Monte Carlo cluster-based permutation approach with 1000 randomisations (Maris & Oostenveld, 2007).

Third, the participants’ performance in each of the auditory tasks was evaluated by means of a mixed model on the ratio of correct responses, by saturation session and difficulty condition, for each of the tasks. This approach was preferred to a repeated measures ANOVA due to missing data points at a subject-wise level (3 on average across tasks).

Fourth, we assessed the efficacy of the auditory tasks in inducing different cognitive load and arousal levels based on their difficulty. Again, given the missing data points, a mixed model was performed with the pupil data of each task as dependent variable. As above, group and session were set as explanatory variables.

Fifth, participants’ subjective evaluation of fatigue was assessed by comparing the values reported before and after the experiment in the Karolinska Sleepiness Scale and Multidimensional Fatigue Inventory. Separate repeated measures ANOVAs were carried out for the responses in the two questionnaires, with session as within-subjects factor and difficulty as between-subjects factor.

Lastly, estimates of behavioural performance change between baseline and conclusion session obtained by the psychometric function, and the change in the both questionnaire scores were correlated. This test was carried out in Matlab.

### Open code and data policy

Source code is available at [https://github.com/ste-ioan/TDT2019], data is available upon request to the corresponding author.

## Results

### Behaviour

The GLMM revealed the quadruple effect of ISI, Saturation, Difficulty and Session to be strongly significant (see Figure 2; X^2^_(2, 46080)_ = 21.96, p < 0.0001; together with Session: X^2^_(2, 46080)_ = 28.18, p < 0.0001; ISI: X^2^_(1, 46080)_ = 95.56, p < 0.0001; ISI and Session interaction: X^2^_(2, 46080)_ = 29.43, p < 0.0001; Saturation, Difficulty and Session interaction: X^2^_(2, 46080)_ = 12.68, p = 0.002; ISI, Difficulty and Session interaction: X^2^_(2, 46080)_ = 51.89, p < 0.0001; and ISI, Saturation and Session interaction: X^2^_(2,46080)_ = 6.88, p = 0.032).

**Figure 2:**
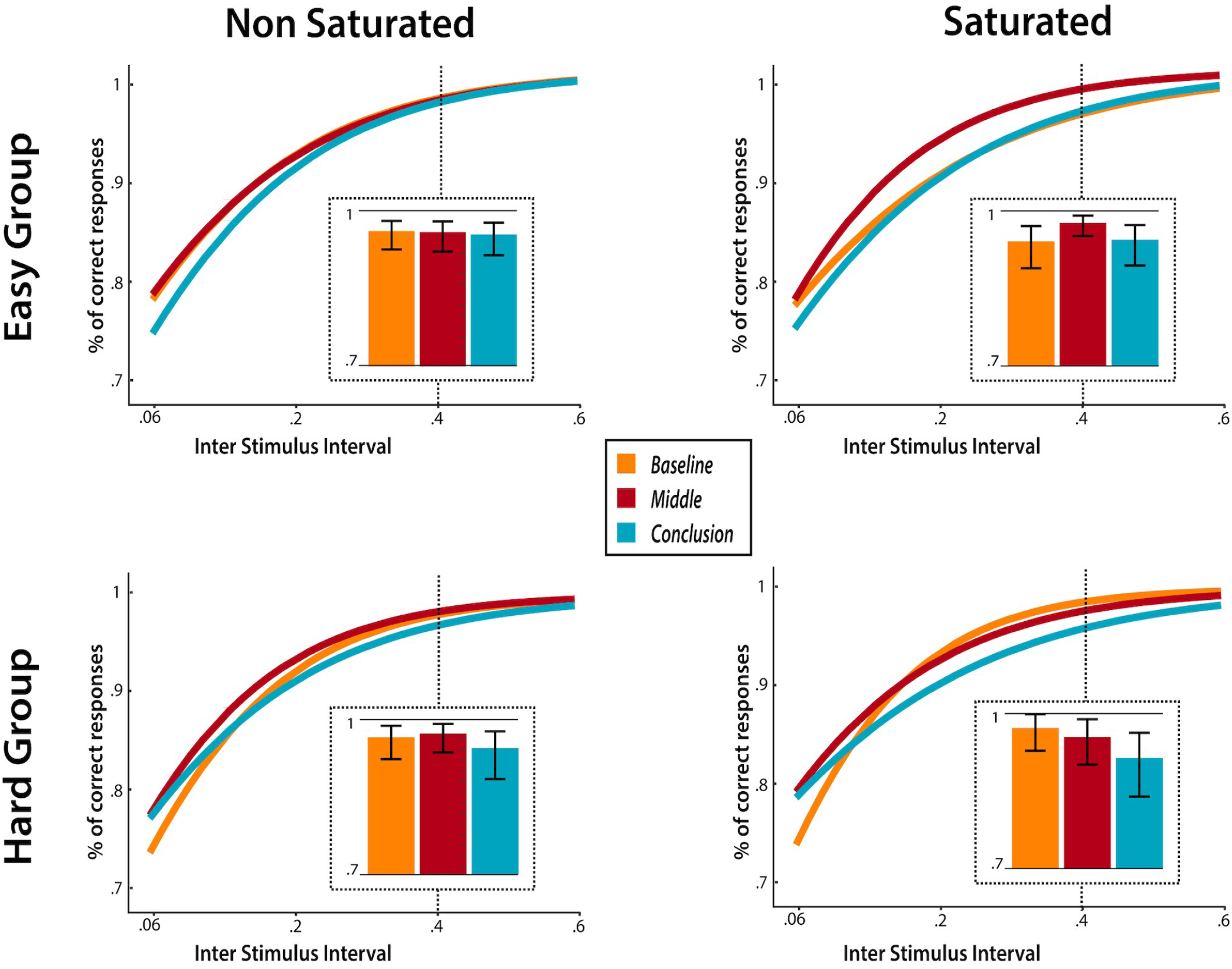
Percent correct responses (y axis), predicted by the GLMM for ISI values ranging from 0.06 to 0.6 (x axis) as a function of saturation condition (left and right columns) and experimental group (top and bottom rows). Error bars depict 95% confidence intervals. Insets: detailed representation of behavioural data for ISI = 400ms.

In order to investigate this four-way interaction, we performed separate GLMMs for the two difficulty groups and for the first and second half of the experiment, applying Holm-Bonferroni correction for multiple comparisons. For the easy group, in the saturated quadrant, performance largely improved between baseline and middle sessions (Z = 3.67, p_corrected_ = 0.001, exp(*β*) = 18.5) and fairly decreased between middle and conclusion sessions (Z = −2.47, p_corrected_ = 0.039, exp(*β*) = 6.9). However, there wasn’t a significant change between baseline and conclusion (Z = 1.06, p_corrected_ = 0.287).

On the other hand, the hard group displayed a major loss of performance in the saturated quadrant, between baseline and middle sessions (Z = −4.168, p_corrected_ = 0.0002, exp(*β*) = 24.4) and baseline and conclusion (Z = −3.93, p_corrected_ = 0.0004, exp(*β*) = 15.3), but not between middle and conclusion (Z = 1.13, p_corrected_ = 0.518), as between these sessions their performance decreased also in the non-saturated quadrant.

To further confirm this finding, we explored its session-wise evolution, by performing the same statistical procedure, per group, session-by-session. The absence of a significant interaction between ISI and Saturation in the baseline session was confirmed, for the easy group (X^2^_(1, 7680)_ = 0.99, p = 0.32), and the hard group (X^2^_(1, 7680)_ = 3.3, p = 0.07). In the middle session, this interaction became significant for both easy (X^2^_(1, 7680)_ = 4.95, p = 0.026) and hard groups (X^2^_(1, 7680)_ = 4.63, p = 0.031) and similarly so in the conclusion session, for easy (X^2^_(1, 7680)_ = 4.87, p = 0.027), and hard (X^2^_(1, 7680)_ = 6.19, p = 0.013).

Thus, repeated visual stimulation led to changes in performance within the saturated quadrant, depending on the difficulty condition. Specifically, participants assigned to the hard condition saw their performance drop, particularly in the first half of the experiment. In contrast, participants in the easy group initially exhibited an improvement in their TDT performance, which disappeared following the second saturation session.

### Electrophysiology

We analysed electrophysiological activity by means of cluster-based permutation methods (Maris & Oostenveld, 2007). We first compared EEG steady-state responses between blocks with stimulation of left and right portion of the visual field in the baseline session across groups (easy, hard). As no clusters were found, we concluded that the side of stimulation did not induce significant differences and grouped left and right stimulation blocks for further analyses. Comparisons of the amplitude of the steady-state response to 7.5Hz stimulation were performed separately for both groups, between baseline and middle, middle and conclusion and between baseline and conclusion. No significant clusters were found.

### Audio task performance

Concerning the performance of the auditory tasks carried out during saturation, the easy group outperformed the hard group in each task. Specifically, in the n-back task the overall mean accuracy of the easy group was 93.3% versus 62.1% of the hard group (t_(45)_ = 10, p < 0.0001). In the side task, the easy group had on average 99.2 % of correct responses, whilst the hard group had an accuracy of 89.1% (t_(46)_ = 3.99, p = 0.0002), Finally, in the pitch-sequence task, the easy group displayed 73.5% average accuracy, while the hard group only reached 49.1% (t_(46)_ = 6.31, p < 0.0001).

### Pupillometry

The analysis of the average pupil size recorded while participants were engaged in the auditory tasks revealed a significant effect of the group condition, in each task (see Figure 3). On top of this, also the effect of session was found to meaningfully impact the pupil size of participants in each task. In the case of the side task, participants in the hard condition had a larger pupil than those in the easy condition (t_(46)_ = 3.94, p = 0.0003) and the whole sample’s pupil size decreased between the first and second saturation sessions (t_(45)_ = −3.58, p = 0.0008). The same result was found also in the n-back task, both in the group-wise effect (t_(46)_ = 2.02, p = 0.049) and in the session-wise effect (t_(45)_ = −2.16, p = 0.036). On the other hand, in the case of the pitch-sequence task, the pupil averages in the easy condition were found to be larger than those in the hard condition (t_(43)_ = − 4.18, p = 0.0001), while the pupil responses decreased in the second session, similarly to the other tasks (t_(40)_ = −2.62, p = 0.012).

**Figure 3:**
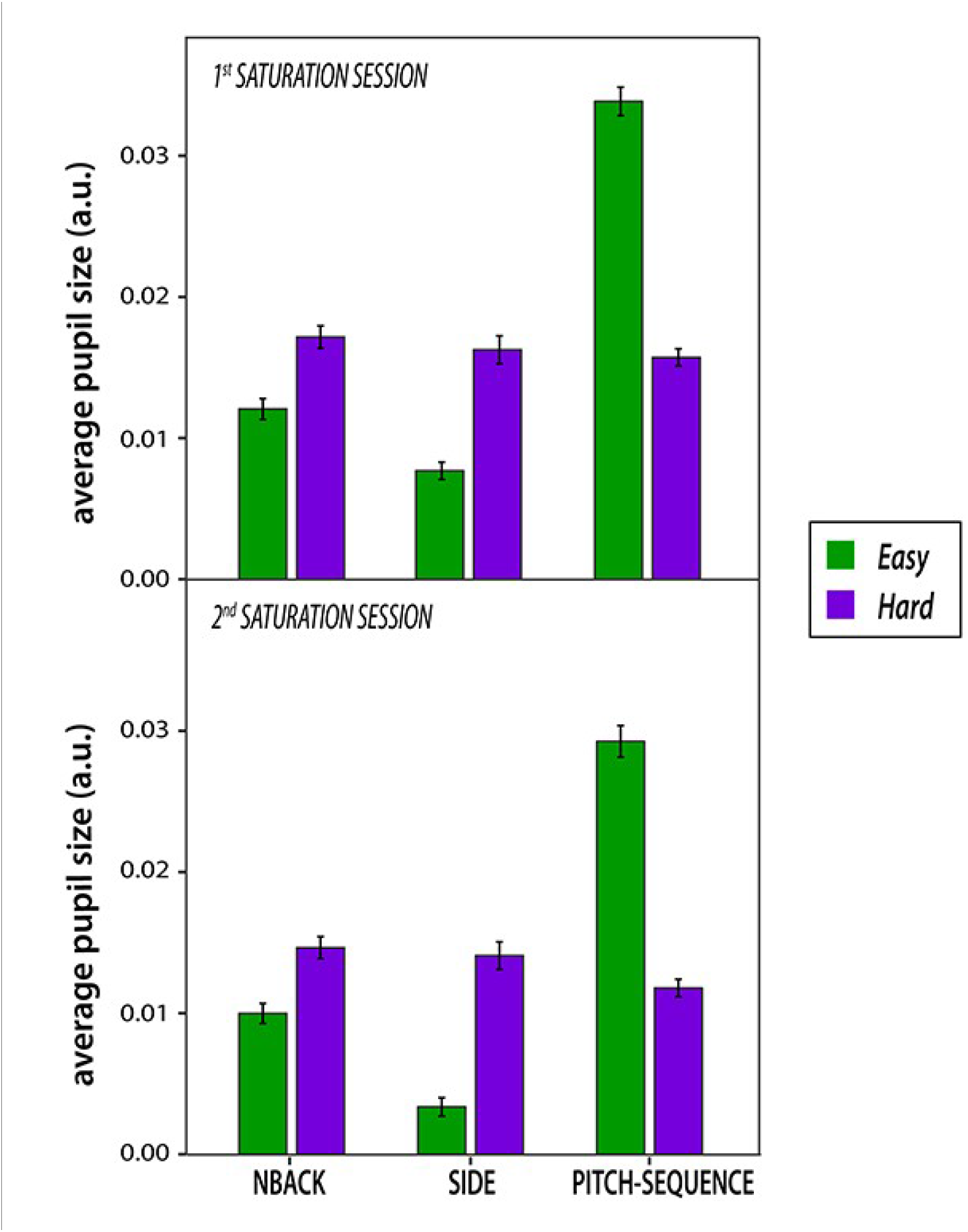
Average pupil size in each auditory task per saturation session, by experimental group

### Self-reported measures of fatigue and sleepiness

The rANOVA on the Multidimensional Fatigue Inventory scores revealed an important increase over the course of the experiment, as highlighted by the significance of the session factor (F_(1, 46)_ = 255.9, p < 0.0001, *η*^2^_p_, while neither the main effect of group, nor their interaction reached significance. Similarly, the Karolinska Sleepiness Scale scores highlighted a strongly significant increase in perceived sleepiness across sessions (F_(1, 46)_ = 176, p < 0.0001, *η* ^2^_p_, while no effect was found for the group condition nor its interaction with session.

Additionally, the increase in subjective fatigue displayed a moderate negative correlation with the change in the behavioural performance (performed across groups given lack of difference shown above; R_(46)_ = −0.29, p = 0.043; see Figure 4). This indicates that larger perceived fatigue related to greater loss in performance, as indexed by behavioural difference between saturated and non-saturated scores across the beginning and conclusion of the experiment. On the other hand, no significant correlation was observed between changes in behaviour and perceived sleepiness (R_S(46)_ = −0.1, p = 0.5)

**Figure 4:**
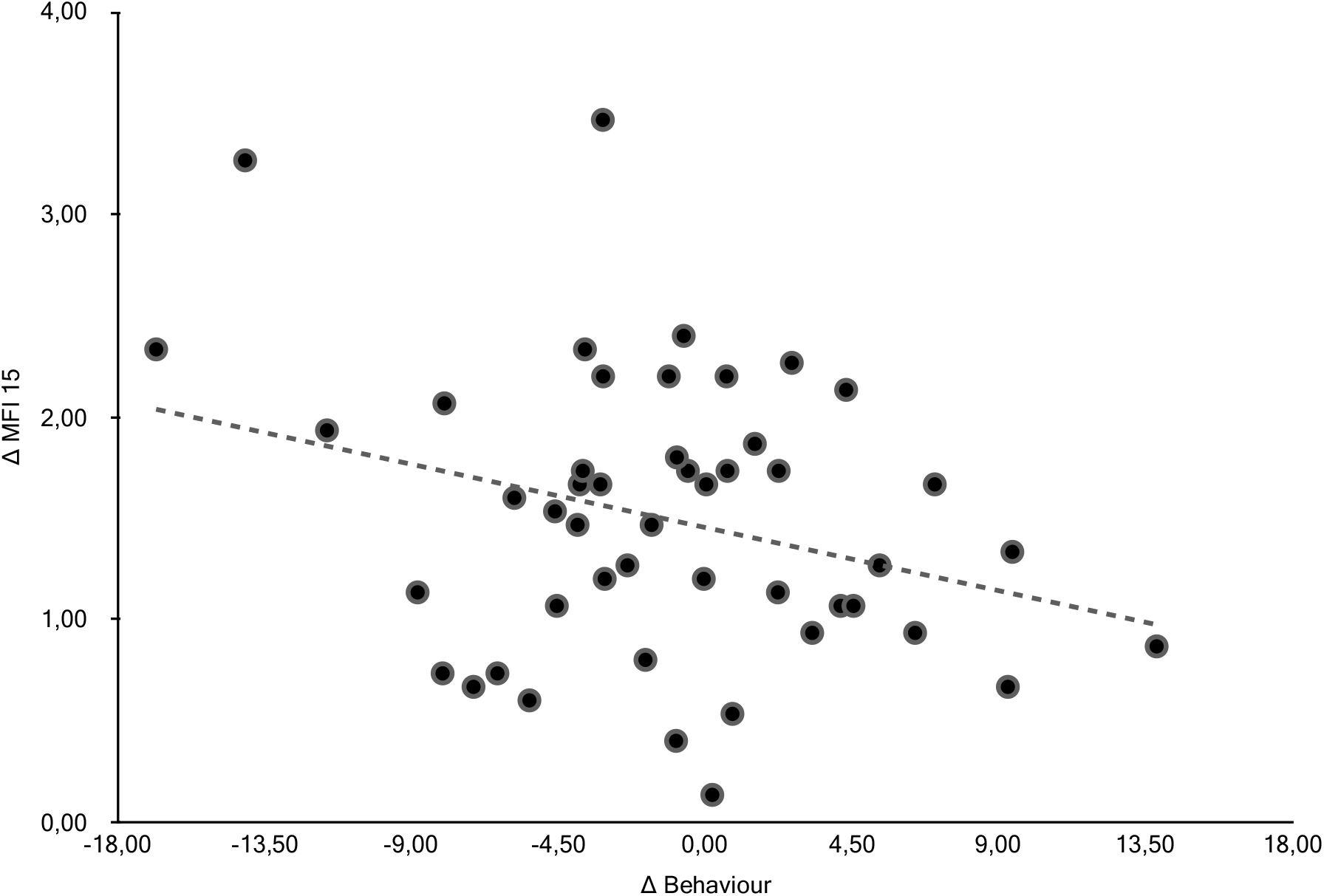
Scatterplot depicting the change in reported fatigue (y axis) in relation to the change in behavioural performance, consisting in the delta between conclusion and baseline in the individual slopes of the participant’s psychometric curves (x axis)

## Discussion

In the present study, we found that prolonged stimulation of a portion of the visual field led to specific deterioration of participants’ performance in that portion of visual space, proving that neural fatigue can be locally and passively induced. Because of its spatial specificity, this neural fatigue phenomenon cannot be explained solely by motivational theories of fatigue. On the contrary, such progressive functional alteration generated by repeated recruitment could constitute one of the mechanisms involved in general fatigue, as suggested by the correlation we observed between the feeling of fatigue and the specific behavioural consequences, which is rarely found in the pertaining literature (DeLuca, 2007).

As predicted, perceptual deterioration was accelerated by concurrent performance of difficult auditory tasks. We argue that increased arousal provoked by the challenging version of the tasks and physiologically indexed by pupil size, led to stronger activations of the simultaneously stimulated visual neurons (Mather et al., 2016; McGinley et al., 2015; Zhang et al., 2020) which in turn caused faster and greater degradation of performance.

Alternatively, it could be argued that the difference in the impact of passive stimulation on performance across groups could be due to cognitive load per se, rather than to the related changes in arousal. Indeed, it cannot be excluded that the effect observed here would require taxing of the cognitive control network (Asplund & Chee, 2013; Blain et al., 2016; Braver, 2012; Mandrick et al., 2013; Niendam et al., 2012) on top of the saturation of the local visual processes. However, it is unclear which interaction mechanism would require the concurrent disruption of spatially specific visual neurons and central cognitive control networks in order to result in local performance decrement. Distinguishing conclusively between our initial hypothesis of arousal modulation amplifying the impact of saturation on the one hand and the possible interaction of global and local saturation effects on the other hand will require more experiments.

Interestingly, on top of perceptual deterioration, we also observed improvement of perceptual performance induced by passive stimulation. This result was restricted to the participants whom underwent the easy versions of the auditory tasks, and faded following the second session of passive stimulation. This observation evokes the passive learning literature, where repeated, passive presentation of visual stimuli leads to ameliorations in subsequent discrimination performance (Watanabe et al., 2001). However, classical passive learning is usually measured on the days following training while, since our experiment was not designed to investigate learning, we did not measure behavioural performance on different days. Therefore, this observation relates better to the older and more general concept of perceptual priming, than to perceptual learning literature (Wiggs & Martin, 1998).

On the other hand, volunteers who were engaged in hard auditory tasks while being visually stimulated failed to show this stimulation-induced enhancement in performance, suggesting that increased neural fatigue induced by their arousal state counteracted perceptual priming. Alternatively, this result may indicate that arousal inhibits perceptual priming processes (though in contradiction of previous accounts; (Thomas & LaBar, 2005)) or that difficult auditory tasks induced diversion of attention away from the visual stimulus (also not in accordance with previous findings (Mulligan, 2003)).

Our results also stand out from previous publications in the field of visual fatigue, which generally rely on self-reported measures of eye discomfort, or other physiological markers such as dry eyes and binocular vision stress (Sheppard & Wolffsohn, 2018). Albeit we did not record these variables, there is no reason to suppose there would be a difference between the experimental groups in any of these measures, as they were exposed exactly to the same visual stimuli under identical conditions and recruited from a homogeneous sample. Furthermore, in the few attempts where this body of literature has sought to assess behavioural effects, it has failed to detect any (Chi & Lin, 1998). Part of our findings are perhaps complementary with the literature on visual adaptation. However, in that case repeated stimulation affects subsequent behaviour on the timescale of seconds (Carandini, 2000), whereas our findings extend over dozens of minutes. Determining with precision how these mechanisms interact would require studying the time course of the effect with better resolution than the one provided in the present study.

From the neurophysiological perspective, we failed to observe significant differences in the amplitude of the electrophysiological brain response to the 7.5 Hz flashing of the stimuli. This lack of effect may rule out the usage of this method as a way to investigate the saturation phenomenon or it may be that our study was not sufficiently powered to detect significant differences in this signal, due to its effect size. It may also indicate that perceptual deterioration originates from decreased signal-to-noise ratio rather than simple decrease in activation magnitude. The use of more spatially sensitive instruments (e.i. fMRI) is warranted to shed light on the brain localisation of the observed behavioural phenomenon, which might be explained by a different factor than electrophysiological magnitude of cortical brain response. For instance, the behavioural changes may be reflected in the neural encoding of the saturating stimuli, and thereby be more aptly investigated by multi-variate pattern analyses (Haxby et al., 2014).

Despite the lack of statistical effects in the recorded brain signal, the fact that the behavioural consequences were influenced by the difficulty of concurrent, cross-sensory auditory tasks still argues in favour of a cortical, as opposed to retinal, origin of the effect. Such cortical origin would be in line with previous perceptual deterioration results, caused by prolonged TDT performance, where decreased primary visual cortex activations were observed (Mednick et al., 2008).

All participants reported an increase in fatigue and sleepiness scores at the end of the experiment, providing evidence for the success of the experimental procedure in impacting the *objective* and *subjective* manifestations of fatigue. In fact, the Multidimensional Fatigue Index results were significantly correlated to the change in saturation-specific behavioural performance. Such correlation cannot be attributable to the behavioural difference between easy and hard groups, since the self-reported measures of fatigue did not significantly differ between them. However, the current design does not allow us to make inferences on the causal direction of this relation.

Analysis of the arousal-related pupil response showed increased dilation in the hard version of the n-back and side tasks, confirming the increased demand associated with these task conditions (van der Wel & van Steenbergen, 2018; Zénon, 2019). However, surprisingly, we found the opposite effect in the pitch-sequence task, the easier group displaying larger average pupil size. This may be due to the excessive demands of this task for those assigned to the hard condition, potentially leading to their disengagement from the task, phenomenon known to reduce pupil size (Hopstaken, van der Linden, et al., 2015). Indeed, the pitch-sequence task had the lowest performance across tasks and conditions (49% of correct responses overall) and several participants spontaneously reported that this task was the most challenging of the three.

In conclusion, we bring new evidence that passive, repeated neuronal activation induces progressive functional alteration under the modulatory influence of cross-sensory cognitive load and arousal: under low cognitive load, we observed an initial perceptual priming effect followed by a decrease to baseline performance, while no changes were detected in the unstimulated portion of visual field. On the other hand, participants under higher cognitive load displayed an early, specific, loss of performance which then generalised to the non-saturated quadrant.

These findings are immune to confounds such as boredom, motivation and level of skill in the fatiguing task (Gergelyfi et al., 2015). Certainly, since all participants carried out the task in the saturated and non-saturated portion of their visual field, any difference in performance between the two, can be solely explained as the by-product of saturation – which was an artifice to keep specific neuronal assemblies active all along.

This neural fatigue phenomenon could justify renewed interest in metabolic accounts of fatigue. Indeed, while the neural mechanism underlying this recruitment-related performance deterioration remains to be investigated, one may speculate the involvement of depletion or accumulation of metabolites (Dalsgaard & Secher, 2007; Dienel & Hertz, 2001; Fairclough & Houston, 2004). The detailed characterisation of such process is matter for subsequent studies, which may disentangle how exactly the phenomena brought forward by the present results, to our best knowledge previously unreported, act and interact with each other.

## References

Åkerstedt, T., Kecklund, G., Alfredsson, L., & Selén, J. (2007). Predicting Long-Term Sickness Absence from Sleep and Fatigue. Journal of Sleep Research, 16, 341–345. https://doi.org/10.1111/j.1365-2869.2007.00609.x

Asplund, C. L., & Chee, M. W. L. (2013). Time-on-task and sleep deprivation effects are evidenced in overlapping brain areas. NeuroImage, 82, 326–335. https://doi.org/10.1016/j.neuroimage.2013.05.119

Aston-Jones, G., & Cohen, J. D. (2005). AN INTEGRATIVE THEORY OF LOCUS COERULEUS-NOREPINEPHRINE FUNCTION: Adaptive Gain and Optimal Performance. Annual Review of Neuroscience, 28(1), 403–450. https://doi.org/10.1146/annurev.neuro.28.061604.135709

Benoit, C.-E., Solopchuk, O., Borragán, G., Carbonnelle, A., Van Durme, S., & Zénon, A. (2019). Cognitive task avoidance correlates with fatigue-induced performance decrement but not with subjective fatigue. Neuropsychologia, 123, 30–40. https://doi.org/10.1016/j.neuropsychologia.2018.06.017

Blain, B., Hollard, G., & Pessiglione, M. (2016). Neural mechanisms underlying the impact of daylong cognitive work on economic decisions. Proceedings of the National Academy of Sciences, 113(25), 6967–6972. https://doi.org/10.1073/pnas.1520527113

Boksem, M. A. S., Meijman, T. F., & Lorist, M. M. (2006). Mental fatigue, motivation and action monitoring. Biological Psychology, 72(2), 123–132. https://doi.org/10.1016/j.biopsycho.2005.08.007

Boksem, M. A. S., & Tops, M. (2008). Mental fatigue: Costs and benefits. Brain Research Reviews, 59(1), 125–139. https://doi.org/10.1016/j.brainresrev.2008.07.001

Borragán, G., Slama, H., Bartolomei, M., & Peigneux, P. (2017). Cognitive fatigue: A Time-based Resource-sharing account. Cortex; a Journal Devoted to the Study of the Nervous System and Behavior, 89, 71–84. https://doi.org/10.1016/j.cortex.2017.01.023

Brainard, D. H. (1997). The Psychophysics Toolbox. Spatial Vision, 10(4), 433–436. https://doi.org/10.1163/156856897X00357

Braver, T. S. (2012). The variable nature of cognitive control: A dual mechanisms framework. Trends in Cognitive Sciences, 16(2), 106–113. https://doi.org/10.1016/j.tics.2011.12.010

Carandini, M. (2000). Visual cortex: Fatigue and adaptation. Current Biology, 10(16), R605–R607. https://doi.org/10.1016/S0960-9822(00)00637-0

Chi, C. F., & Lin, F. T. (1998). A comparison of seven visual fatigue assessment techniques in three data-acquisition VDT tasks. Human Factors, 40(4), 577–590. https://doi.org/10.1518/001872098779649247

Dalsgaard, M. K., & Secher, N. H. (2007). The brain at work: A cerebral metabolic manifestation of central fatigue? Journal of Neuroscience Research, 85(15), 3334–3339. https://doi.org/10.1002/jnr.21274

DeLuca, J. (2007). Fatigue as a Window to the Brain. MIT Press.

Dienel, G. A., & Hertz, L. (2001). Glucose and lactate metabolism during brain activation. Journal of Neuroscience Research, 66(5), 824–838. https://doi.org/10.1002/jnr.10079

Dongen, H. P. A. V., Belenky, G., & Krueger, J. M. (2011). Investigating the temporal dynamics and underlying mechanisms of cognitive fatigue. https://doi.org/10.1037/12343-006

Drugowitsch, J. (2019). Variational Bayesian inference for linear and logistic regression. ArXiv:1310.5438 [Stat]. http://arxiv.org/abs/1310.5438

Fairclough, S. H., & Houston, K. (2004). A metabolic measure of mental effort. Biological Psychology, 66(2), 177–190. https://doi.org/10.1016/j.biopsycho.2003.10.001

Gallucci, M. (2019). GAMLj: General analyses for linear models. https://gamlj.github.io/

Gergelyfi, M., Jacob, B., Olivier, E., & Zénon, A. (2015). Dissociation between mental fatigue and motivational state during prolonged mental activity. Frontiers in Behavioral Neuroscience, 9. https://doi.org/10.3389/fnbeh.2015.00176

Goode, J. H. (2003). Are pilots at risk of accidents due to fatigue? Journal of Safety Research, 34(3), 309–313. https://doi.org/10.1016/S0022-4375(03)00033-1

Grillon, C., Quispe-Escudero, D., Mathur, A., & Ernst, M. (2015). Mental fatigue impairs emotion regulation. Emotion (Washington, D.C.), 15(3), 383–389. https://doi.org/10.1037/emo0000058

Haxby, J. V., Connolly, A. C., & Guntupalli, J. S. (2014). Decoding Neural Representational Spaces Using Multivariate Pattern Analysis. Annual Review of Neuroscience, 37(1), 435–456. https://doi.org/10.1146/annurev-neuro-062012-170325

Hockey, R. (2013, May). The Psychology of Fatigue: Work, Effort and Control. Cambridge Core. https://doi.org/10.1017/CBO9781139015394

Hopstaken, J. F., Linden, D. van der Bakker, A. B., & Kompier, M. A. J. (2015). A multifaceted investigation of the link between mental fatigue and task disengagement. Psychophysiology, 52(3), 305–315. https://doi.org/10.1111/psyp.12339

Hopstaken, J. F., van der Linden, D., Bakker, A. B., & Kompier, M. A. J. (2015). The window of my eyes: Task disengagement and mental fatigue covary with pupil dynamics. Biological Psychology, 110, 100–106. https://doi.org/10.1016/j.biopsycho.2015.06.013

Jones, B. E. (2003). Arousal systems. Frontiers in Bioscience, 8(6), s438–451. https://doi.org/10.2741/1074

Karni, A., & Sagi, D. (1991). Where practice makes perfect in texture discrimination: Evidence for primary visual cortex plasticity. Proceedings of the National Academy of Sciences of the United States of America, 88(11), 4966–4970. https://doi.org/10.1073/pnas.88.11.4966

Kontsevich, L. L., & Tyler, C. W. (1999). Bayesian adaptive estimation of psychometric slope and threshold. Vision Research, 39(16), 2729–2737. https://doi.org/10.1016/S0042-6989(98)00285-5

Kurzban, R., Duckworth, A., Kable, J. W., & Myers, J. (2013). An opportunity cost model of subjective effort and task performance. The Behavioral and Brain Sciences, 36(6). https://doi.org/10.1017/S0140525X12003196

Mandrick, K., Derosiere, G., Dray, G., Coulon, D., Micallef, J.-P., & Perrey, S. (2013). Prefrontal cortex activity during motor tasks with additional mental load requiring attentional demand: A near-infrared spectroscopy study. Neuroscience Research, 76(3), 156–162. https://doi.org/10.1016/j.neures.2013.04.006

Maris, E., & Oostenveld, R. (2007). Nonparametric statistical testing of EEG- and MEG-data. Journal of Neuroscience Methods, 164(1), 177–190. https://doi.org/10.1016/j.jneumeth.2007.03.024

Mather, M., Clewett, D., Sakaki, M., & Harley, C. W. (2016). Norepinephrine ignites local hot spots of neuronal excitation: How arousal amplifies selectivity in perception and memory. The Behavioral and Brain Sciences, 39, e200. https://doi.org/10.1017/S0140525X15000667

McGinley, M. J., David, S. V., & McCormick, D. A. (2015). Cortical membrane potential signature of optimal states for sensory signal detection. Neuron, 87(1), 179–192. https://doi.org/10.1016/j.neuron.2015.05.038

Mednick, S. C., Drummond, S. P. A., Arman, A. C., & Boynton, G. M. (2008). Perceptual Deterioration is Reflected in the Neural Response: Fmri Study of Nappers and Non-Nappers. Perception, 37(7), 1086–1097. https://doi.org/10.1068/p5998

Mulligan, N. W. (2003). Effects of cross-modal and intramodal division of attention on perceptual implicit memory. Journal of Experimental Psychology. Learning, Memory, and Cognition, 29(2), 262–276. https://doi.org/10.1037/0278-7393.29.2.262

Niendam, T. A., Laird, A. R., Ray, K. L., Dean, Y. M., Glahn, D. C., & Carter, C. S. (2012). Meta-analytic evidence for a superordinate cognitive control network subserving diverse executive functions. Cognitive, Affective, & Behavioral Neuroscience, 12(2), 241–268. https://doi.org/10.3758/s13415-011-0083-5

Norcia, A. M., Appelbaum, L. G., Ales, J. M., Cottereau, B. R., & Rossion, B. (2015). The steady-state visual evoked potential in vision research: A review. Journal of Vision, 15(6), 4–4. https://doi.org/10.1167/15.6.4

Oostenveld, R., Fries, P., Maris, E., & Schoffelen, J.-M. (2011). FieldTrip: Open Source Software for Advanced Analysis of MEG, EEG, and Invasive Electrophysiological Data [Research Article]. Computational Intelligence and Neuroscience; Hindawi. https://doi.org/10.1155/2011/156869

Pelli, D. G. (1997). The VideoToolbox software for visual psychophysics: Transforming numbers into movies. Spatial Vision, 10(4), 437–442.

Raslear, T. G., Hursh, S. R., & Van Dongen, H. P. A. (2011). Chapter 10—Predicting cognitive impairment and accident risk. In H.P.A. Van Dongen & G. A. Kerkhof (Eds.), Progress in Brain Research (Vol. 190, pp. 155–167). Elsevier. https://doi.org/10.1016/B978-0-444-53817-8.00010-4

Ricci, J. A., Chee, E., Lorandeau, A. L., & Berger, J. (2007). Fatigue in the U.S. workforce: Prevalence and implications for lost productive work time. Journal of Occupational and Environmental Medicine, 49(1), 1–10. https://doi.org/10.1097/01.jom.0000249782.60321.2a

Schellekens, J. M. H., Sijtsma, G. J., Vegter, E., & Meijman, T. F. (2000). Immediate and delayed after-effects of long lasting mentally demanding work. Biological Psychology, 53(1), 37–56. https://doi.org/10.1016/S0301-0511(00)00039-9

Sheppard, A. L., & Wolffsohn, J. S. (2018). Digital eye strain: Prevalence, measurement and amelioration. BMJ Open Ophthalmology, 3(1), e000146. https://doi.org/10.1136/bmjophth-2018-000146

Swaen, G. M. H., Amelsvoort, L.G.P.M. van Bültmann, U., & Kant, I. J. (2003). Fatigue as a risk factor for being injured in an occupational accident: Results from the Maastricht Cohort Study. Occupational and Environmental Medicine, 60(suppl 1), i88–i92. https://doi.org/10.1136/oem.60.suppl_1.i88

The jamovi project. (2019). Jamovi (1.1) [Computer software]. Error! Hyperlink reference not valid.

Thomas, L., & LaBar, K. (2005). Emotional arousal enhances word repetition priming. Cognition and Emotion, 19(7), 1027–1047. https://doi.org/10.1080/02699930500172440

van der Wel, P., & van Steenbergen, H. (2018). Pupil dilation as an index of effort in cognitive control tasks: A review. Psychonomic Bulletin & Review, 25(6), 2005–2015. https://doi.org/10.3758/s13423-018-1432-y

Wang, C.-A., Baird, T., Huang, J., Coutinho, J. D., Brien, D. C., & Munoz, D. P. (2018). Arousal Effects on Pupil Size, Heart Rate, and Skin Conductance in an Emotional Face Task. Frontiers in Neurology, 9, 1029. https://doi.org/10.3389/fneur.2018.01029

Watanabe, T., Náñez, J. E., & Sasaki, Y. (2001). Perceptual learning without perception. Nature, 413(6858), 844–848. https://doi.org/10.1038/35101601

Wiggs, C. L., & Martin, A. (1998). Properties and mechanisms of perceptual priming. Current Opinion in Neurobiology, 8(2), 227–233. https://doi.org/10.1016/S0959-4388(98)80144-X

Zénon, A. (2019). Eye pupil signals information gain. Proceedings of the Royal Society B: Biological Sciences, 286(1911), 20191593. https://doi.org/10.1098/rspb.2019.1593

Zénon, A., Solopchuk, O., & Pezzulo, G. (2019). An information-theoretic perspective on the costs of cognition. Neuropsychologia, 123, 5–18. https://doi.org/10.1016/j.neuropsychologia.2018.09.013

Zhang, D., Yan, X., She, L., Wen, Y., & Poo, M. (2020). Global enhancement of cortical excitability following coactivation of large neuronal populations. Proceedings of the National Academy of Sciences, 117(33), 20254–20264. https://doi.org/10.1073/pnas.1914869117

